# Longitudinal characterization of compound action potentials in chronic vagus nerve recordings in mice

**DOI:** 10.1101/2025.07.21.665755

**Authors:** Shubham Debnath, Ibrahim T. Mughrabi, Todd J. Levy, Fylaktis Fylaktou, Nilay Kumar, Yousef Al-Abed, Stavros Zanos, Theodoros P. Zanos

## Abstract

The vagus nerve (VN) mediates bidirectional communication between the body and brain to maintain physiological homeostasis; likewise, alterations in ongoing vagal signaling may be indicators of disease and/or contribute to disease pathogenesis. Even though extensively documented in acute experiments, ongoing vagal activity has not been characterized longitudinally, over days or weeks, in mice, a preferred preclinical model. In addition, even though many VN recordings in mice occur during anesthesia, the effects of anesthesia on vagal signaling are unknown. This study uses a chronic implant mouse model to record vagal activity in anesthetized and awake, behaving animals for an average of 10 weeks and up to 6 months. Individual compound action potentials (CAPs) are tracked across multiple days by quantifying comparisons in features, including firing rates, waveform shape, inter-CAP interval histograms, and phase-locking to cardiac and respiratory signals while demonstrating long-term electrode-nerve interface viability and stable signal-to-noise ratios. Additionally, cytokine challenge experiments produced detectable CAP responses up to 3 months after electrode implantation. Lastly, awake recordings incorporated video analysis to identify and remove motion artifacts to preserve and extract neural and cardiac recordings during behavior. Results reveal diverse CAP populations with diverse physiological coupling and firing rates modulated by anesthesia. This work highlights the potential of chronic VN recordings to assess long-term changes in vagal activity in health and disease, with implications in discovery of autonomic markers of disease and closed-loop VNS stimulation strategies.

## Introduction

The autonomic nervous system monitors physiological changes and responds to maintain internal homeostasis. In particular, the vagus nerve (VN) plays a key role in the control of homeostasis, as it takes part in several autonomic reflexes [1–4]. The role of autonomic reflexes in regulating inflammation through inflammatory reflexes has received attention, as inflammation contributes to several common chronic diseases [5–9]. The emerging field of bioelectronic medicine has focused on modulating inflammatory reflexes via neurostimulation to treat chronic diseases [10–12]. However, less attention has been given towards resolving spontaneous activity of autonomic reflexes in general, and vagal activity in particular, even though altered vagal signaling may be a marker of disease, and may also be contributing to disease pathogenesis.

Studies have linked afferent (sensory) fiber activity in the VN to inflammatory, metabolic, and cardiopulmonary physiological functions. For example, sensory signals in anesthetized mice have been linked to the presence of cytokines [13–15] and to hypoglycemic states [16]. Studies in anesthetized rats linked vagus electroneurogram with cardiovascular function related to epileptic seizures [17], classifying firing patterns related to breathing and blood glucose levels [18], and distinguishing motor and sensory signals during upper airway obstruction [19]. In most of these studies, recording from autonomic nerves is done in rodent models; however, the small size of rodent VN poses technical and surgical challenges for recording, especially in chronic experiments [20,21]. In the rat model, recordings have been made possible in freely moving animals to measure spike patterns related to locomotor and peripheral organ activity [22] and classifying behaviors like eating [23], while chronic implants have allowed vagal recordings for up to 11 weeks after implantation relating to heart rate variability, vagal tone, and circadian rhythms [23–27]. Cardiac and respiratory modulation has also been recorded from the human vagus nerve using ultrasound-guided microneurography [28–29]. While these studies are important for clinical translation and understanding VN’s role in homeostasis, no chronic recording studies with longitudinal data (anesthetized or awake) in the same animal have been performed in the mouse, the most common preclinical model for disease pathophysiology, genetic screenings, and pharmacological mechanisms [30,31]. Additionally, the effect of anesthesia on VN activity has not been documented in mice, even though it can affect electrophysiological measurements of nervous activity and other experimental parameters [32–36].

To utilize sensory signals to diagnose emerging conditions or evaluate treatment efficacy, VN signals must be reliably recorded chronically and decoded to identify biomarkers related to inflammation or markers of autonomic dysfunction. A chronically implanted mouse model has been developed and shown to deliver functional stimulation for at least four weeks after cuff implantation, in anesthetized [37] and more recently in awake mice [38]. The same chronic VN implant was used to record VN activity from implanted cuff electrodes in mice, along with simultaneously recorded electrocardiogram (ECG) and nasal air flow to monitor heart and respiratory activity, respectively, while modulating anesthesia level, injecting cytokines, and during awake light phase behavior. VN activity was recorded for up to six months in anesthetized animals, while tracking neural signal-to-noise ratio (SNR) to quantify implant stability and heart rate threshold (HRT) current intensity to confirm cuff-nerve interface viability. Additionally, compound action potentials (CAPs) were quantified and several of their features were documented, including phase-locking to cardiac and respiratory signals, to track unique CAPs across recording sessions. VN activity was also recorded from awake, behaving animals while tracking movement to remove epochs related to motion-related artifacts and extract neural signals. Our chronic VN recording model in mice can be used to document changes in VN activity in health and in disease states.

## Methods

### Electrode preparation

Two types of cuff electrodes were commercially fabricated and used in this study (Figure 1A). MicroLeads150 μm cuff electrodes (MicroLeads Neuro, Somerville, MA) utilize a self-closing mechanism and are constructed using silicone, polymide, and platinum iridium (Figure 1A, left), while CorTec 100 μm microsling cuffs (CorTec, Freiburg, Germany) utilize a buckle-closing mechanism and are constructed using silicone, platinum iridium, and a parylene C coating (Figure 1A, middle). Lead wires to trimmed to 2.3 to 2.5 cm and soldered to gold socket pins (Figure 1A, right). Impedance was measured in saline for each electrode at 1 kHz using a MicroProbes tester (MicroProbes for Life Science, Gaithersburg, MD), and soldered electrodes were sterilized before implantation by submerging in 0.55% ortho-phthaladehyde (Cidex OPA, Advanced Sterilization Products, Irvine, CA) solution for 15 minutes and rinsed with sterile saline.

**Figure 1.**
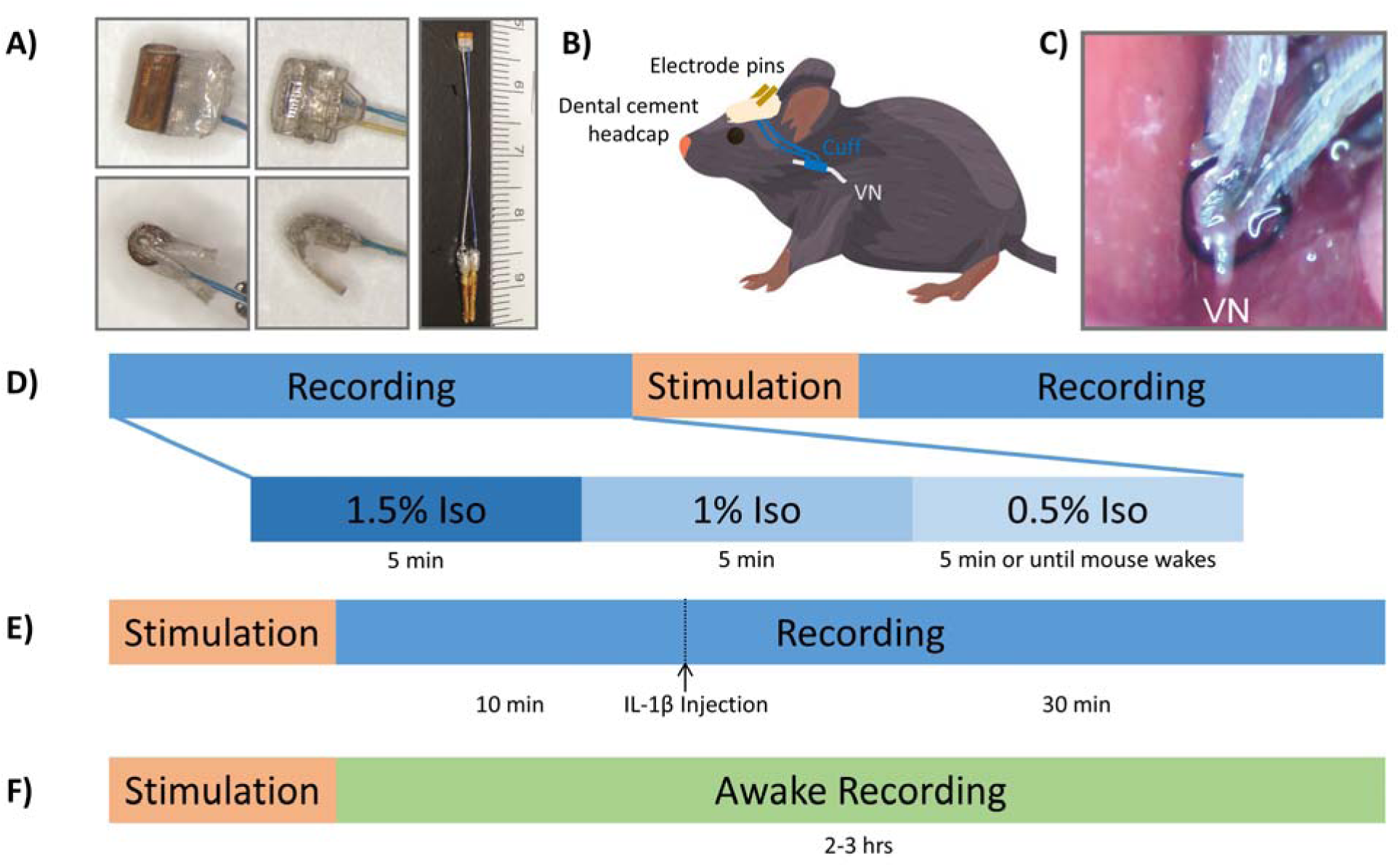
Electrodes implantation and experiments. (A) Shown are front (top panels) and side views (bottom panels) of cuff electrodes used in chronic implants produced by MicroLeads (left panels) or CorTec (middle panels). Lead wires were cut to 2.5-3.0 cm and soldered to gold pins (right panel). Adapted from [37]. (B, C) The cuff was placed on the left cervical VN, with lead wires and gold pins guided and secured to a dental cement headcap on exposed skull. Adapted from [37]. (D) Anesthetized recording sessions consisted of two recording blocks before and after a short stimulation block, which was only to establish the HRT and confirm a functional neural interface. Each recording block consisted of 5 minutes each at 1.5%, 1.0%, and 0.5% isoflurane, in succession. (E) Injection recording sessions were also under anesthesia. A short stimulation block was done, again only to establish HRT and confirm a functional interface, followed by 10 minutes of baseline recording. Upon IL-1β injection, data were collected for a further 30 minutes. (F) For awake recordings, stimulation under anesthesia was used to establish HRT, and then neural data were recorded for two to three hours.

### Surgical procedure

All animal care, surgical, and research procedures were performed in accordance with relevant ethical guidelines and approved by the Institutional Animal Care and Use Committee (IACUC) at the Feinstein Institutes for Medical Research, protocol number 2019-010. Male C57BL/6 mice were housed under 12 hr light/dark cycle with ad libitum access to water and food. Cuff electrodes were implanted on the left cervical VN, and lead wires were tunneled to a dental cement anchored headcap on the skull (Figure 1B), with slight adjustments from surgical methods described by [37]. The surgical procedure took place under isoflurane anesthesia (4% for induction, 1.5-2% for maintenance) with regular warm saline supplemented intraoperatively, starting the animal in a supine position to shave the neck area with an electric shaver and disinfect the skin using betadine and 70% alcohol. Once cleaned, the animal was turned to a prone position and placed in a stereotaxic frame to stabilize the head and access the skull. The head hair was also shaved and trimmed with scissors then the scalp was disinfected with betadine and 70% alcohol. A local analgesic (bupivacaine) was applied before cutting skin over the center of the skull to expose bregma and lambda. The skull was cleaned with three to four alternating rinses of hydrogen peroxide and saline, while scoring the skull with 18g needle between rinses. The skull was then dried using compressed air, and acrylic dental cement (Metabond, Parkell, Edgewood, NY) was applied to the right side and allowed to set and cure. A subcutaneous tunnel was created between the animal’s eye and ear by blunt dissection down the skull incision towards the left neck. The mouse was then flipped back to supine position and placed on a heated surgical platform. The neck area was once cleaned again with betadine and 70% ethanol, and a 1-cm vertical incision was made about 0.5 cm left of midline, starting at the sternal notch. The parotid gland was displaced away from subcutaneous tissue, and muscles in the anterior triangle were shifted to expose the carotid sheath; the VN was carefully bluntly dissected away from the carotid artery. The cuff electrode to be implanted was tunneled via the pre-formed subcutaneous tunnel from the skull incision to the neck incision site using fine straight forceps. The VN was lifted and placed into the cuff, the closing mechanism was put in place, and then the VN was placed back close to its original anatomic position (Figure 1C). A brief current stimulus was delivered via the gold pins to measure a heart rate response and confirm electrode placement and interface viability. Once established, the neck incision was closed with 6-0 nylon suture and the animal was again placed in a prone position. Dental cement was again applied to anchor the external gold pins and liquid adhesive was applied to seal the skin-cement junction. Surgically implanted mice were placed in single housed cages and monitored under conscious and mobile, treated with meloxicam (5 mg/kg), and then allowed to recover for 3 days before any experiments.

For awake recording experiments, animals were additionally implanted with three platinum iridium wires, placed through the subcutaneous neck tunnel and tethered to the chest wall to record ECG [38]. Both the cuff and ECG wires were connected to a multi-channel nano-connector (Omnetics Connector Corporation, Minneapolis, MN) and cemented to the skull using dental cement, as described earlier.

### Vagus nerve recording experiments

VN activity was recorded from implanted cuff electrode in two experiments under anesthesia, while simultaneously recording ECG and nasal air flow to monitor heart and breathing rates, respectively. ECG was recorded from needle electrodes placed in the animal’s forelimbs with ground at the animal’s left hindlimb, while nasal air flow was measured by placing a thermocouple wire into the animal’s nostril. For the first experiment (Figure 1D), each session involved a recording block before and after VN stimulation to confirm viability of the electrode-nerve interface by establishing the HRT needed to induce a heart rate change, or at least a 20% drop from baseline heart rate. Each recording block consisted of 5-minute periods at each of three levels of isoflurane anesthesia (1.5%, 1%, and 0.5% of 1 L/min O_2_). These sessions were repeated daily in the same animal until HRT could not be safely determined or mechanical failure. The second experiment (Figure 1E), no more than once per week, was performed under anesthesia. After again confirming the electrode-nerve interface via stimulation and baseline recording for 10 minutes, VN activity was recorded for 30 minutes after an IL-1β (35 ng/kg) injection. For awake recording experiments (Figure 1F), stimulation was applied under anesthesia to establish an HRT and confirm the interface viability, followed by awake, behaving recordings of two to three hours.

### Signal decomposition and compound action potential feature extraction

Raw neurogram was recorded from 2 channels at a sample rate of 30 kHz before being band-pass filtered from 100 to 4 kHz and notch comb filtered at the first five multiples of 60 Hz (Figure 2, top). An impedance matching approach was used to remove common-mode interference from the two electrode contacts, including ECG and respiratory artifacts, to extract and retain neural signals [39]. From the cleaned neural signals, individual CAPs were extracted by adaptive thresholding, and a previously described framework was applied to cluster groups of waveforms [15]. Dimensionality reduction used a t-distributed stochastic neighbor embedding (t-SNE) method to visualize, and unsupervised CAP clustering utilized a density-based spatial clustering of applications with noise (DBSCAN) algorithm to determine the number of waveform groups and assign individually extracted CAPs to those groups. Meanwhile, raw physiology, including ECG and respiration, were recorded at a sampling rate of 1 kHz, producing cardiac and respiratory rates and phase (Figure 2, bottom). Phase was assigned by defining 0 as peak inhalation and R-peak for respiratory and cardiac signals, respectively, with -π defined as half-way to the previous peak and π as half-way to the next peak. After defining CAP groups from neural data and cardiac and respiratory phase from physiological data, each group of CAPs could be described by a number of features, namely average waveform shape, firing rates and changes due to anesthesia level or cytokine injection (in anesthetized studies), inter-CAP interval histograms, and phase-locking histograms on respiratory and cardiac cycles (Figure 2, bottom right).

**Figure 2.**
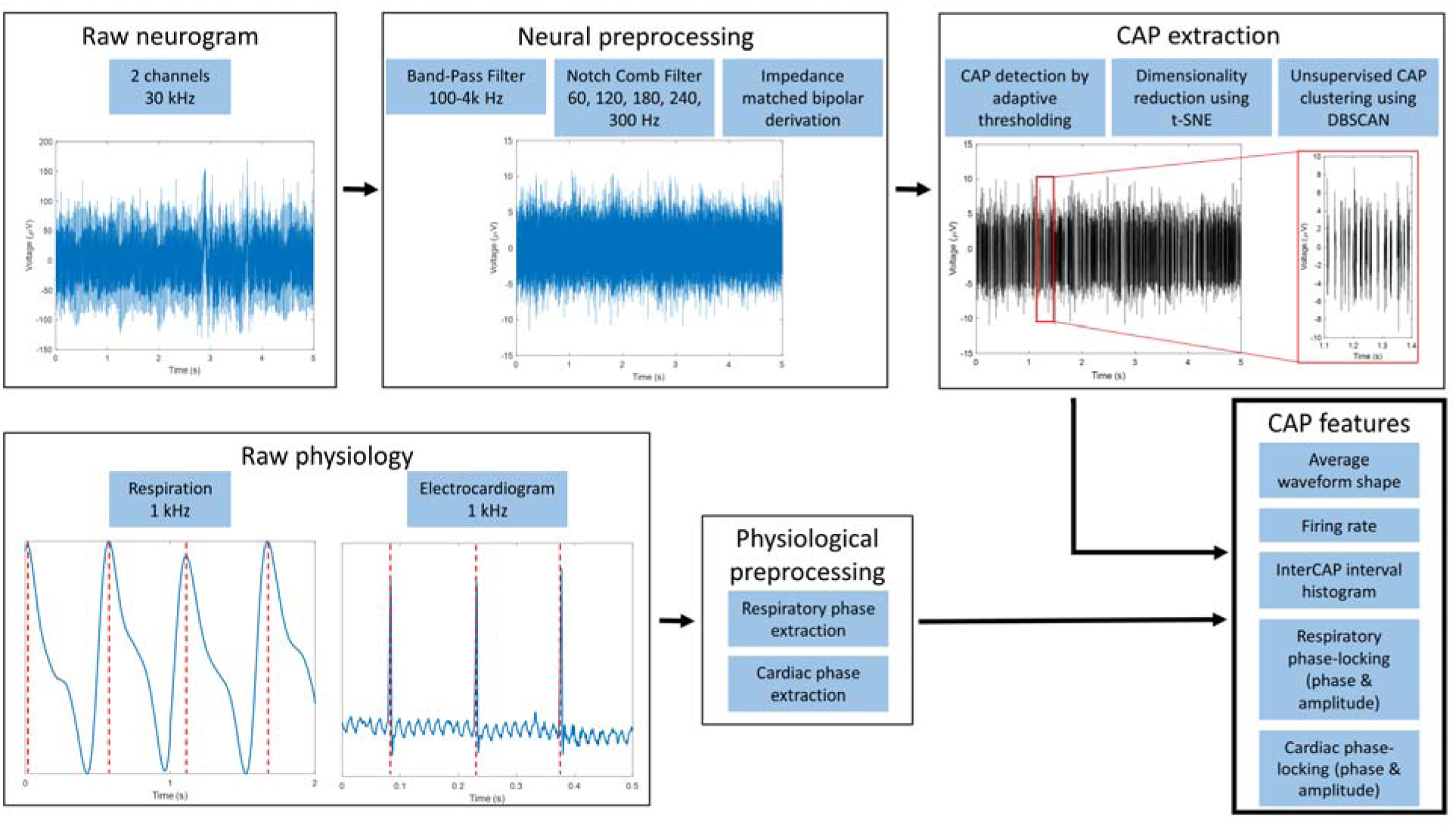
Analysis pipeline and feature extraction. The raw neurogram (top row) is recorded from 2 channels at a sampling frequency of 30 kHz. This signal is preprocessed by bandpass and notch comb filters, along with a impedance matching algorithm used to remove cardiac and respiratory-related noise. Compound action potentials are isolated by a series of steps including adaptive thresholding, dimensionality reduction, and unsupervised clustering. CAP features extracted are average waveform shape, firing rate, and interCAP intervals. The raw physiology includes respiration and electrocardiogram. Phase is assigned by defining 0 as peak inhalation and R-peak for the respiratory and cardiac signal, respectively (denoted by vertical red dotted lines). The respiratory and cardiac phase-locking characteristics for each CAP was also extracted as a feature.

### Similarity index calculation and longitudinal signal tracking

To track unique CAPs over multiple recording days or sessions in the anesthetized experiments, these five distinct CAP features, and their changes relative to variation in the rate of isoflurane, were compared day-to-day. For each animal and extracted CAPs on consecutive recording days, cross-correlation coefficients were calculated between average waveform shape, firing rate over time relative to isoflurane rate, inter-CAP interval histograms, and phase-locking with respiration and cardiac activity relative to isoflurane rate (Figure 3A-E); the five calculated cross-correlation values were averaged to calculate a similarity index between CAPs extracted on different days (Figure 3F). A threshold value to determine whether CAPs sustained over multiple days was found by comparing the similarity index of CAPs from different animals, such that they are definitively not the same CAPs, and setting a cutoff at the 95% percentile. CAPs recorded on consecutive days with a composite similarity index > 0.8 were determined to likely be the same unique CAP and their duration of continuous detection was tracked to determine the stability of vagus signals.

**Figure 3.**
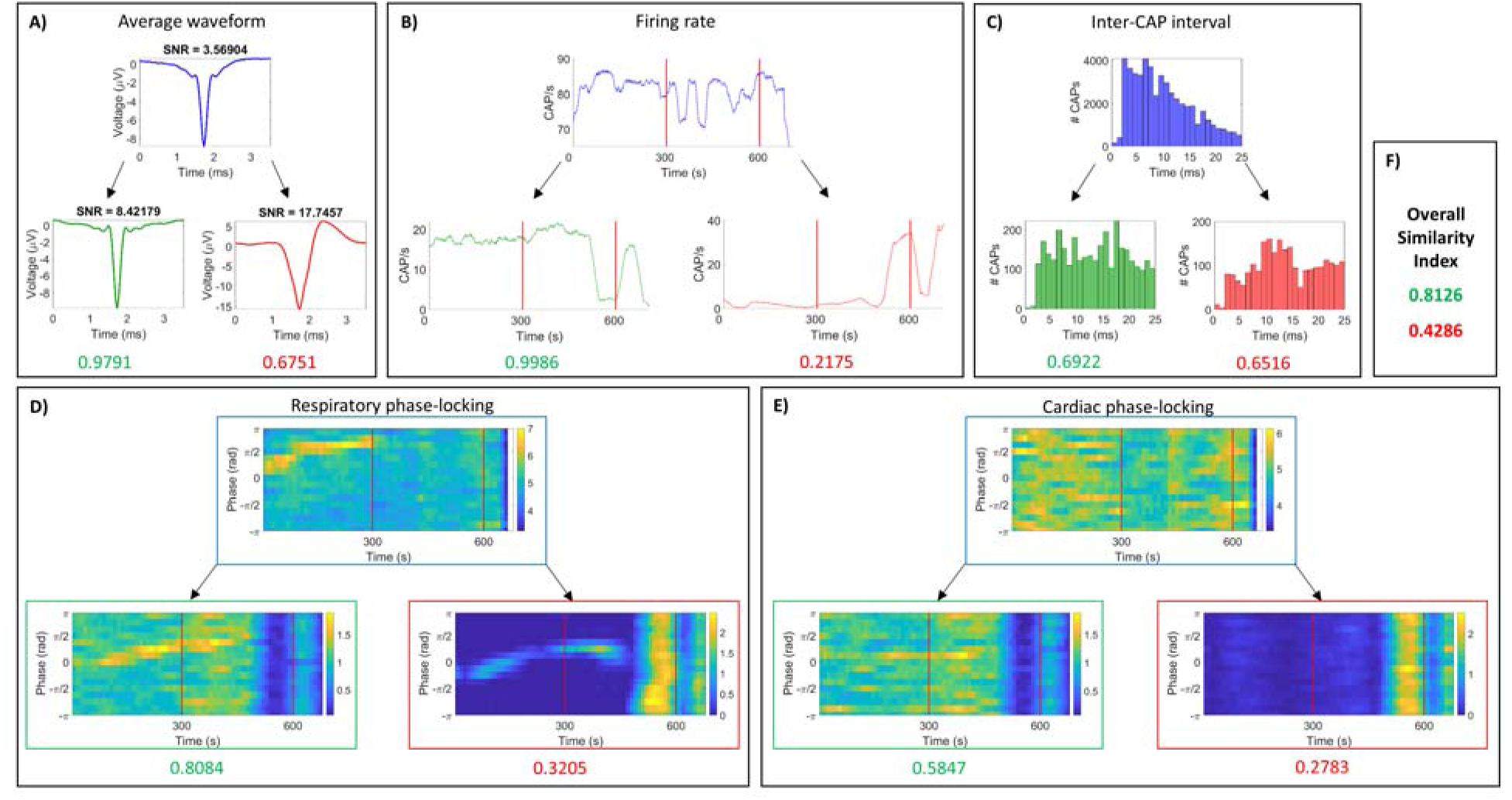
Example of CAP features and similarity index calculation. Cross-correlation coefficients were calculated for five distinct CAP features and their changes relative to changes in rate of isoflurane. Shown is a breakdown of these comparative values between CAPs that have a high similarity index (blue and green) and CAPs that have a low similarity index (blue and red). The green scores in each panel are the cross-correlation coefficient between blue and green CAP features, while the red scores are the cross-correlation coefficient between blue and red CAP features. The average of values for (A) average waveform, (B) firing rate over time and isoflurane rate, (C) inter-CAP interval histogram, (D) phase-locking with respiration relative to isoflurane rate, and (E) phase-locking with cardiac activity relative to isoflurane rate are shown in (F) as the overall similarity index.

### Awake recordings and movement epoch rejection

For awake, behaving recordings, mice were placed in a custom cage made up of a clear, acrylic cylinder (15.24 cm in diameter, 20.32 cm in height) with a lid consisting of an integrated commutator system (PI Technologies, Roanoke, VA) connected to the Neurochip 3 stimulating/recording system [40]. Mice were connected to the commutator system using a flexible cable that allowed free movement and behavior (Figure 4A). Neural recordings, sampled at 30 kHz, took place during the animal’s light phase to reduce movement, as mice perform more sustained activity during a dark phase [41], and animals were filmed to extract movement times that may lead to excess noise in neural recordings. For body part tracking, DeepLabCut, a transfer learning algorithm with deep neural networks, was used to estimate pose by tracking body parts in 2D video without additional markers [42,43]. Videos of five individual animals were cropped from two separate recordings of multiple animals. Frames were extracted at two different time windows towards the beginning and end of each animal’s video. A total of 20 frames per video were automatically selected using the “kmeans” algorithm within the “automatic” method of the “extract_frames” function. Training data was generated by labelling the following components in the mice: (i) nose, (ii) headcap, (iii-iv) left and right ears, (v-vi) two points at the tail-base, and (vii-viii) two points at the end of the tail. 95% of the total data was used for training. A ResNet-50-based neural network was applied with default parameters for a maximum of 20000 training iterations. The network was validated, with a test error of 8.57 pixels and train error of 3.24 pixels (cropped image size: 268 x 268). After applying a p-cutoff of 0.6, the conditioned train error was 3.26 pixels, and the conditioned test error was 3.85 pixels. This network was then used to analyze videos from similar experimental settings. Each labeled video was further qualitatively assessed for the quality of tracking. The position and velocity of these labels (Figure 4B) were used to determine epochs of animal movement that could be discarded in the synced cardiac (Figure 4C) and neural signals (Figure 4D). Firing rates for extracted CAPs (Figure 4E, F) could also be extracted and aligned with the velocity signal to determine the effects of movement on extracted values.

**Figure 4.**
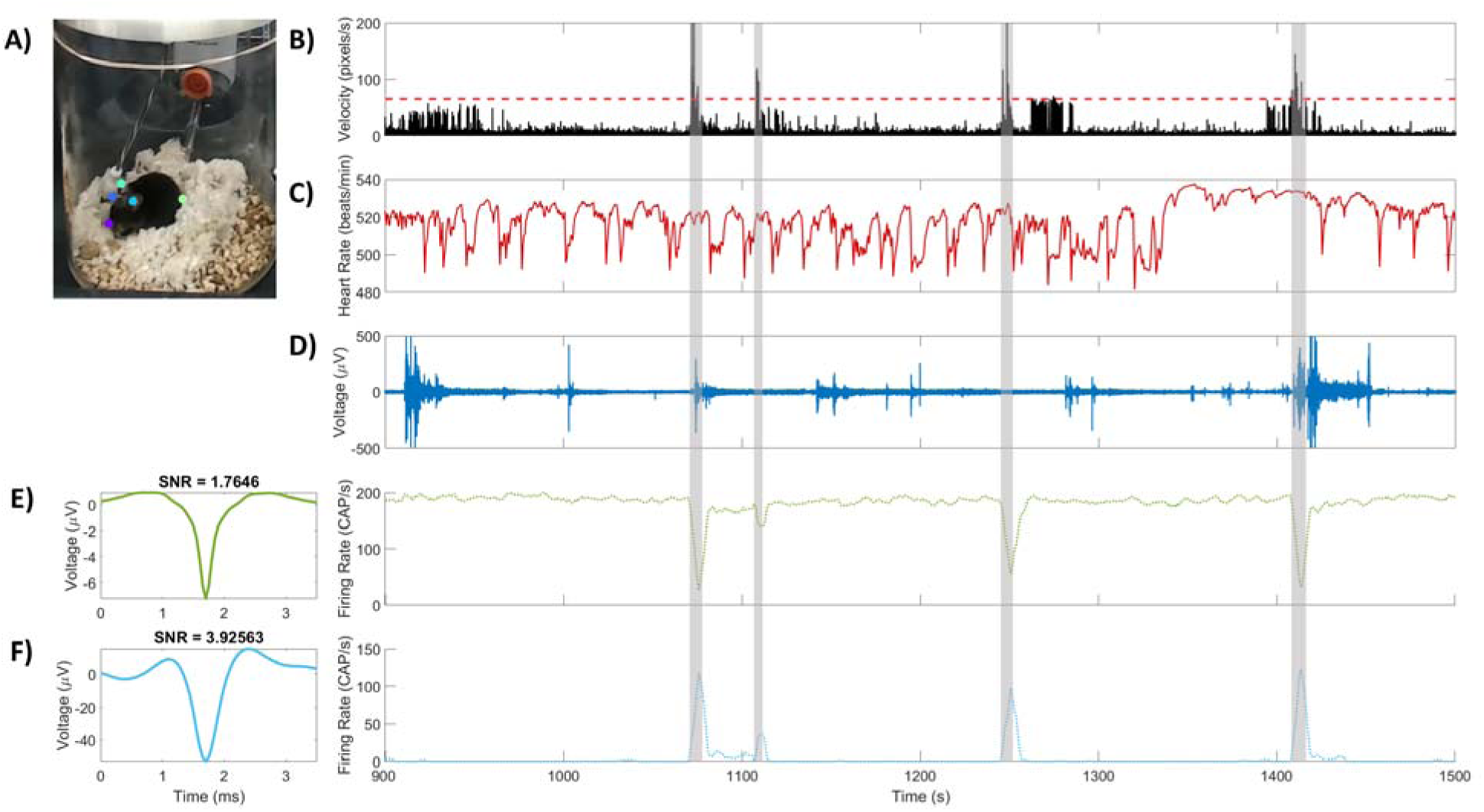
Example of 10 minutes of recording in an awake animal. (A) A screenshot from video produced by DeepLabCut shows markers at the nose, headcap, right ear, left ear, and base of tail tracked by the software. Continuous position of these markers was tracked, and overall velocity was used to track movement (B). A threshold of 70 pixels/second (red horizontal line) was determined to signify a moving animal based on recorded video (translucent grey epochs). (C) Heart rate of the animal was also recorded during awake recordings, with corresponding and time-synced neurogram (D). From this neurogram, two CAPs were extracted shown by their average waveform, SNR, and firing rate (E, F). Drops in firing rate in E (green) and increases in firing rate in F (light blue) correspond to threshold crossing in mouse velocity (B). Additionally, not all movements relate to significant changes in the neurogram.

## Results

### Yield of neural recordings

Anesthetized recordings were completed daily, for 12 to 185 days after surgical implant (Table 1). One group of animals (A-F) were euthanized at 18-19 days; a second group of implanted animals (G-Q) remained viable for recording under anesthesia for an average of 70 days. Of those, 5 implants failed due to wire, electrode, or headcap damage, all within the first 3 weeks post-implantation. The remaining 11 animals were viable for an average of 115.5 days. A total of 575 CAPs were extracted from a total of 23 animals (A-Q), and of these, 339 were unique among other CAPs extracted from the same animals, based on a similarity score. Cytokine injections were performed in 2 animals (P and Q).

**Table 1.** Summary of animals for awake and anesthetized experiments. Shown are all animals used for anesthetized and awake recordings, along with their implant durations, number of recording sessions, electrode type, and extracted CAPs. In the implant duration column, ^ denotes implants ended due to use in another study, while * signifies implant end due to mechanical failure by wire, electrode, or headcap damage. Animals in bold italics were also used for cytokine injection experiments, with further information in Table 3.

| Animal | Recorded State | Implant Duration (d) | Recording Sessions | Electrode | Total Extracted CAPs | Unique CAPs |
| --- | --- | --- | --- | --- | --- | --- |
| A | Anesthetized | 18^ | 3 | ML | 4 | 2 |
| B | Anesthetized | 18^ | 3 | ML | 5 | 3 |
| C | Anesthetized | 18^ | 3 | ML | 5 | 4 |
| D | Anesthetized | 19^ | 3 | ML | 5 | 3 |
| E | Anesthetized | 19^ | 5 | ML | 5 | 4 |
| F | Anesthetized | 19^ | 3 | ML | 3 | 1 |
| G | Anesthetized | 12* | 6 | CT | 13 | 8 |
| H | Anesthetized | 68 | 38 | CT | 86 | 48 |
| I | Anesthetized | 24 | 13 | CT | 34 | 22 |
| J | Anesthetized | 12* | 4 | CT | 8 | 6 |
| K | Anesthetized | 185 | 117 | CT | 120 | 66 |
| L | Anesthetized | 18* | 5 | CT | 10 | 9 |
| M | Anesthetized | 21* | 7 | CT | 18 | 12 |
| N | Anesthetized | 113 | 64 | CT | 74 | 45 |
| O | Anesthetized | 19* | 9 | CT | 20 | 14 |
| ***P*** | Anesthetized | 147 | 79 | CT | 88 | 58 |
| ***Q*** | Anesthetized | 156 | 87 | CT | 50 | 34 |
| R | Awake | 20 | 1 | CT | 3 |  |
| S | Awake | 20 | 1 | CT | 2 |  |
| T | Awake | 20 | 1 | CT | 4 |  |
| U | Awake | 31 | 2 | CT | 5 |  |
| V | Awake | 31 | 2 | CT | 7 |  |
| W | Awake | 31 | 2 | CT | 6 |  |

Awake recordings were collected 1 or 2 times in 6 animals (R-W), up to 1 month after electrode cuff implantation; 27 CAPs were extracted in total in awake sessions, after removing movement epochs. An example of movement epochs (translucent grey vertical bars) can be seen in 10 minutes of awake recording in Figure 4, with a threshold of 70 pixels/second (Figure 4B, red horizontal line) determined to signify a moving animal based on recorded video. Heart rate and synced neurogram were also recorded during the awake recordings (Figure 4C-D). From this neurogram, two CAPs were extracted; drops in firing rate in 4E and increases in firing rate in 4F correspond to threshold crossings in mouse velocity. Not all movement periods related to significant changes in the raw neurogram despite the changes in firing rate.

### CAP features in longitudinal recordings

CAP features over implant lifetime can be seen in Figure 5. Over 27 weeks of recording, the average SNR over all extracted CAPs was 9.47, and remained relatively stable, until week 25 at which point it worsened (Figure 5A). Meanwhile, the HRT remained below 1500 μA, though it progressively increased over time (Figure 5B). Currents below 1000 μA were able to elicit a minimum of 20% drop in heart rate for over 20 weeks after implant, indicating a well-functioning neural implant. Lastly, the lifetime of unique CAPs was tracked based on a composite similarity score > 0.8 in consecutive recording days or sessions. In CAPs captured in multiple sessions, firing rates and phase-locking characteristics with respiratory or cardiac cycles were maintained, along with similar inter-CAP histograms and waveform shape, as shown in the example in Figure 3 (green). Most CAPs (71%) only were captured in a single session, while 12% were stable for more than 3 days (Figure 5C).

**Figure 5.**
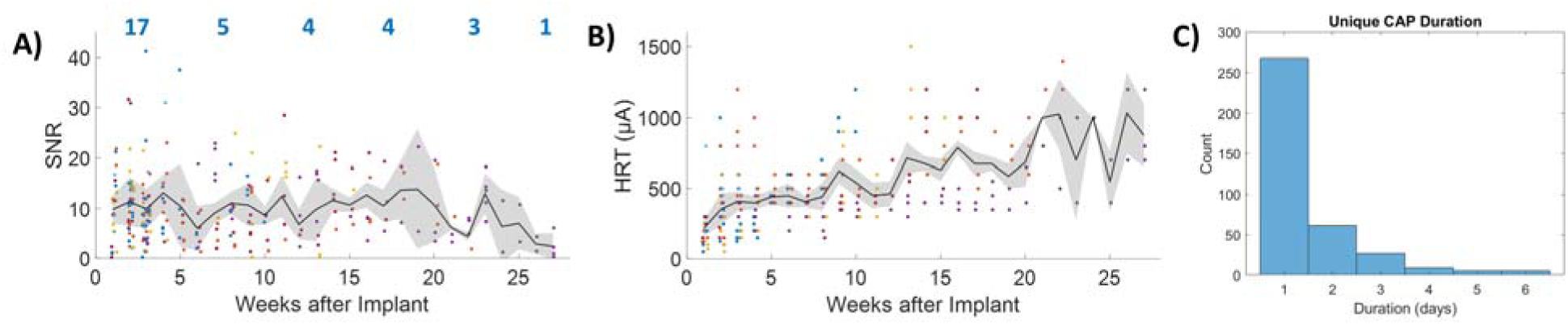
CAP features in longitudinal recordings. (A) Mean SNR calculated from recordings that occurred during a given week post-implant (black), and standard deviation of SNR, across recording and animals (grey envelope). Individual CAP SNRs are shown as colored dots, with each color representing a different animal. The number of animals represented by the traces in these panels decreases as implants fail, shown by blue values above plot that represent the number of animals in each 5-week period. Raw durations per implant are shown in Table 1. (B) The mean (black) and standard deviation (grey envelope) of HRTs increases over time, and implants were considered failed if no response was detected upon stimulation up to 2000 μA. Individual HRTs are shown as colored dots, with each color representing a different animal. (C) Histogram of lifetimes of stable, unique CAPs, in days. Most CAPs were detected for only one day, while 12% were stable for more than 3 consecutive days.

### Effect of level of anesthesia on CAP features

Average firing rates of unique CAPs were calculated at different levels of anesthesia, from 1.5% to 1% to 0.5% isoflurane; CAP firing rates were also calculated during awake conditions (denoted as 0% isoflurane). Most CAPs had firing rates below 25 aps (action potentials per second), with rates during awake generally higher than during anesthesia (Figure 6A); in contrast, the probability of CAPs with firing rates over 100 aps decreased at lower levels of anesthesia (Figure 6A). Based on their FR at the highest level of anesthesia (1.5% isoflurane; FR_1.5_), 3 groups of CAPs emerge. CAPs in the first group, with low FR_1.5_ (FR_1.5_ ≤ 15 aps), increase in firing rate with lowered anesthesia, with a significant increase from 1.0% to 0.5% isoflurane (Figure 6B; blue traces). CAPs in the second group, with intermediate FR_1.5_ (FR_1.5_ = 15-80 aps) maintain relatively stable FR, with no significant changes at different levels of anesthesia (Figure 6B; red traces). Finally, CAPs in the third group, with high FR_1.5_ (FR_1.5_ = 80-150 CAP/s), show significant drops in FR at lighter levels of anesthesia (Figure 6B; yellow traces). Firing rates for CAPs detected during awake recordings were either > 100 or < 40 CAP/s (Figure 6B, green points).

**Figure 6.**
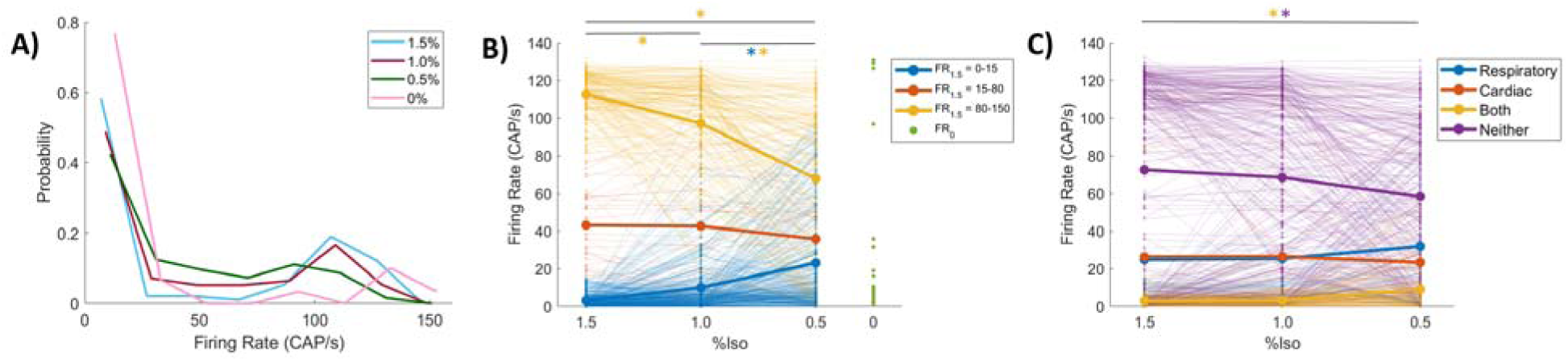
CAP firing rates at different levels of anesthesia. (A) Probability density function of all CAP firing rates at each of 1.5%, 1.0%, and 0.5% isoflurane, as well as the awake recordings (0%). (B) Firing rate changes vs rate of isoflurane, where each trace represents a single extracted CAP, while the colors signify 3 categories based on firing rate at 1.5% isoflurane (FR_1.5_), including 0-15, 15-80, 80-150 CAPs per second. The average of each category is shown by the bolder line of each color. The green points show firing rates of all CAPs detected during awake recordings (FR_0_). (C) Again, firing rate changes vs rate of isoflurane, where each trace represents a single extracted CAP, but colors here represent function determined by significant phase-locking with respiratory signal, cardiac signal, both, or neither. The average of each category is shown by the bolder line of each color. For all figures, * signifies *p* < 0.05.

The same CAPs were also analyzed with regard to their associated physiological characteristics. Examples of CAPs associated with specific phases (phase-locked) in the respiratory cycle, cardiac cycle, or both are shown in Supplementary Figures 1-3. CAPs phase-locked to either the respiratory or cardiac cycle did not significantly change with anesthesia. However, CAPs with no phase-locking and CAPs that showed significant phase-locking with both cardiac and respiratory signals significantly decreased and increased, respectively, at lower levels of anesthesia (Figure 6C). CAPs with no locking had high FR_1.5_, while CAPs that locked with both signals had very low FR_1.5_ (Figure 6C).

CAP phase-locking features and their firing rates can be further classified in relation to anesthesia level. The number of CAPs extracted for each range of FR_1.5_ and phase-locking characteristics are shown in Table 2. Figure 7 shows only CAPs with significant respiratory and/or cardiac phase-locking. The strength of phase-locking and preferred phase can be visualized with rate of anesthesia. For respiratory phase-locking, the amplitude for Group 1 CAPs was significantly higher at 1.5% isoflurane, while amplitude for Group 2 CAPs was significantly lower at 0.5% isoflurane. No Group 3 CAPs showed significant phase-locking with respiratory signals. For cardiac phase-locking, amplitude for Group 1 CAPs was significantly higher at 1.5%. The amplitude of cardiac phase-locking was significantly higher for Group 2 CAPs at 0.5%. There were no significant trends in preferred phase related to isoflurane rate change, but all CAPs average preferred phase was near 0, or peak inhalation and at the R-peak in the ECG signal.

**Figure 7.**
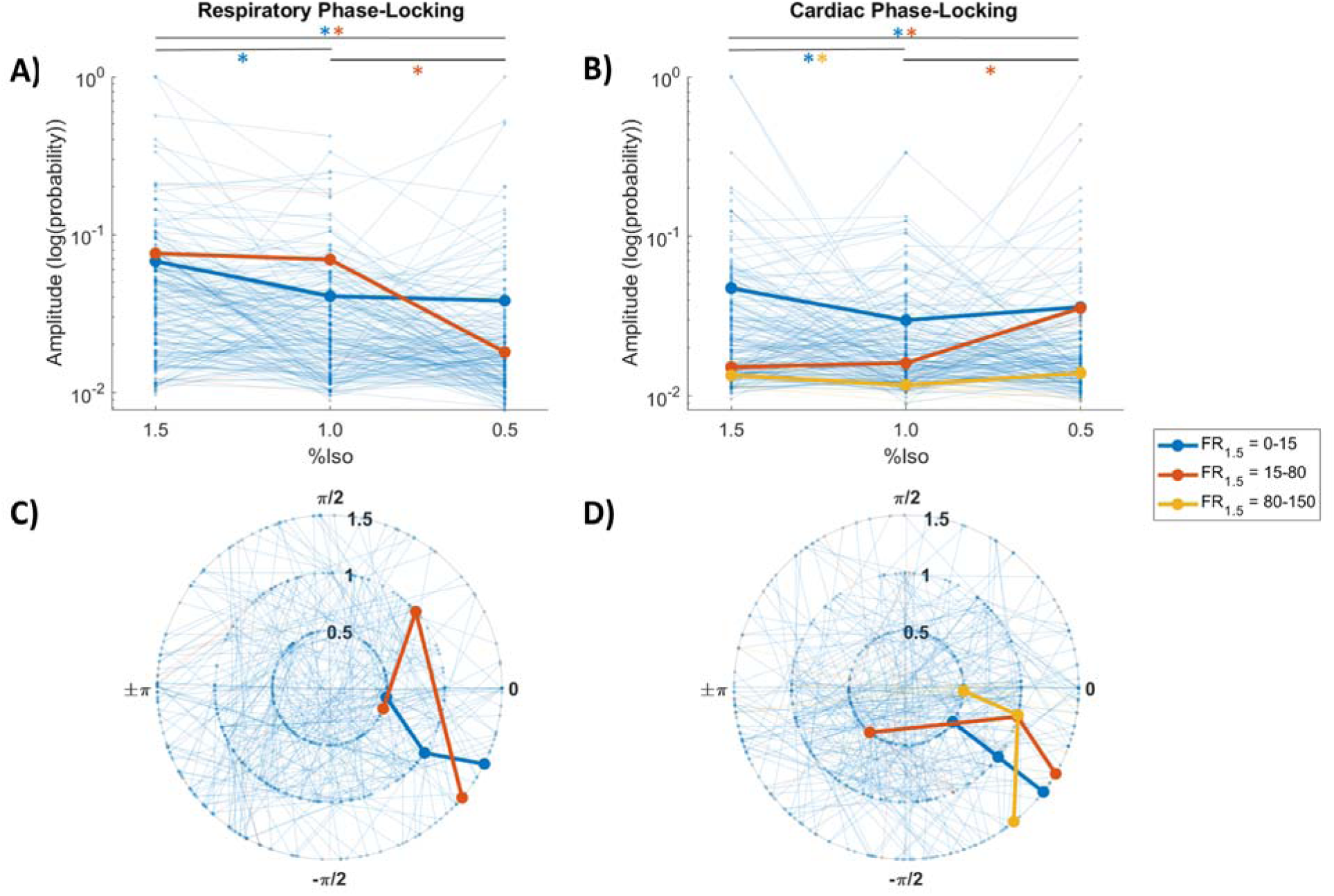
CAP phase-locking features vs rate of isoflurane. Only CAPs with significant respiratory or cardiac (or both) phase-locking are included in these plots. (A,C) Respiratory phase-locking amplitude and phase. (B,D) Cardiac phase-locking amplitude and phase. Each trace represents a single extracted CAP, with colors defined by FR_1.5_, including 0-15, 15-80, 80-150 CAPs per second. No CAPS with FR_1.5_ > 80 showed significant respiratory phase-locking. The average of each category is shown by the bolder line of each color. For all figures, * signifies *p* < 0.05.

**Table 2.** Summary of CAP phase-locking characteristics. Shown are the number of extracted CAPs that display phase-locking with respiratory signal, cardiac signal, both, or neither.

|  | FR<sub>1.5</sub> = 0-15 | FR<sub>1.5</sub> = 15-80 | FR<sub>1.5</sub> = 80-150 | Totals |
| --- | --- | --- | --- | --- |
| No Phase-Locking | 108 | 29 | 191 | **328** |
| Respiratory Phase-Locking Only | 47 | 8 | 0 | **55** |
| Cardiac Phase-Locking Only | 33 | 4 | 10 | **47** |
| Cardiac & Respiratory Phase-Locking | 144 | 1 | 0 | **145** |
| **Totals** | **332** | **42** | **201** | **575** |

### Effect of cytokine challenges on CAP features

Cytokine injection experiments were performed in n = 2 animals and beyond three months after electrode cuff implantation (Table 3). Three injections were administered in each animal, with a total of 16 extracted CAPs. Of these, 7 responder CAPs (43.75%) were observed to increase in firing rate after injection. An example of a responder CAP shown in Figure 8, with extracted CAP features like average waveform and inter-CAP interval (Figure 8A and D, respectively). The vertical red line corresponds to cytokine injection time, and there are no clear changes in heart rate or respiratory rate (Figure 8C). However, firing rate for this CAP increased within 2 minutes (Figure 8B), with significant phase-locking with the respiratory signal (Figure 8E, F). The average respiratory waveform does not alter after the cytokine injection, showing that the breathing activity itself does not change (Figure 8G). There is no significant cardiac phase-locking (Figure 8H, I), but the R-R interval histogram shows a bimodal distribution after injection (Figure 8J). This CAP stops firing after approximately 25 minutes after injection.

**Figure 8.**
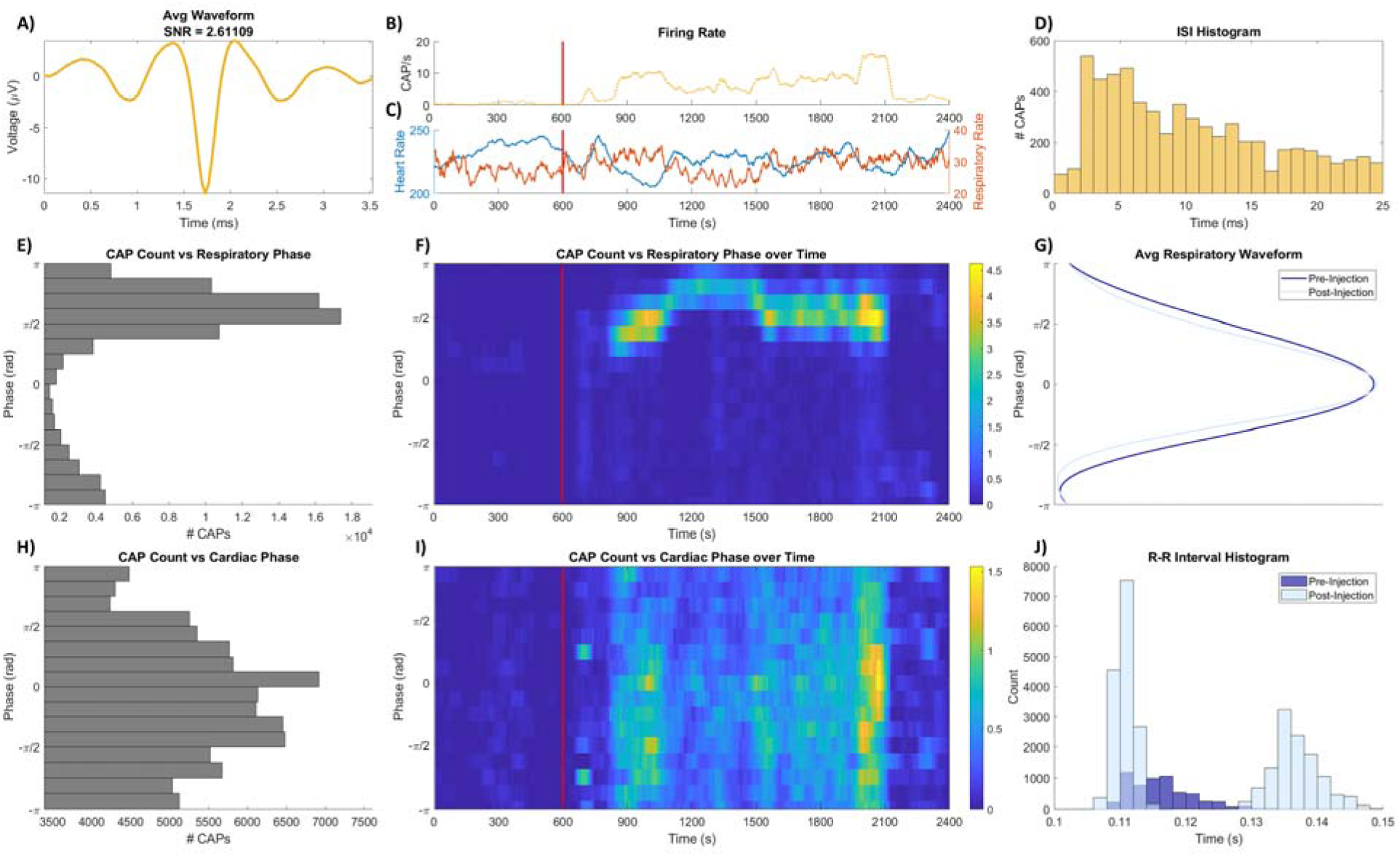
Example of CAP and physiological features during IL-1β injection. Shown is an example of an extracted CAP and its features during baseline and after cytokine injection. Vertical red lines in the middle panels correspond to the time of injection. (A) Average waveform and SNR. (B) Firing rate over the recording session, with (C) corresponding heart (blue) and respiratory rate (red) of the animal. (D) Inter-spike interval histogram. (E,F) CAP count vs respiratory phase, where 0 represents peak inhale, and –π and π represent halfway between the previous and next inhale, respectively. (G) Average shape of respiratory waveform pre- and post-injection. (H,I) CAP count vs cardiac phase, where 0 represents peak of the R complex, and –π and π represent halfway between the previous and next R complex, respectively. (J) Histogram of R-R intervals pre- and post-injection.

**Table 3.** Summary of animals in cytokine injection experiments. Shown are the animals that received cytokine injections, along with the number of injections received (no more than one per week), extracted CAPs, and those that responded to injection by an increase in firing rate after injection.

| Animal | Number of Injections | Total Extracted CAPs | Responders |
| --- | --- | --- | --- |
| P | 3 | 7 | 3 |
| Q | 3 | 9 | 4 |

## Discussion

This study presents a chronic recording model implemented to record CAPs from the mouse VN for over 6 months in both anesthetized and awake animals, with stable SNRs and HRTs. Firing rates and phase-locking characteristics of CAPs were quantified relative to level of anesthesia; with decreasing anesthesia, Group 1 CAPs increase in firing rate and decrease in physiological coupling, Group 2 CAPs’ firing rates remain steady but shift in coupling based on related physiological signal, and Group 3 CAPs decrease in firing rate. A longitudinal analysis of CAP recordings in the same animal is possible with this model by using a quantitative index to determine whether a unique CAP is recorded in consecutive days. Previous literature in this area has focused on reporting electrophysiological recordings from anesthetized terminal experiments in multiple preclinical animal models, while any chronic and/or awake recordings have been observed in rat models.

In this work, the cuff electrode-nerve interface remained viable for over 26 weeks post-implantation, confirmed by tracking the HRT current. Removing implants that were ended due to use in another study, implanted animals were recorded from for an average of 10 weeks; removing implants that failed due to wire breakage or headcap damage, 6 implants lasted an average of 16.5 weeks. Additionally, the SNR of recorded and extracted CAPs remained stable over this implant period. Previous studies utilizing a cuff electrode in mice have only been for terminal experiments [14–16] or for chronic stimulation [36,37]. Chronic studies in rats have utilized intraneural electrodes out to 11 weeks, with higher SNR and individual spiking activity, as expected when recording from within the nerve [44]. No previous study has characterized stability of recorded CAPs from the vagus nerve. By measuring cross-correlations between extracted CAP features and their physiological relationship with other biological signals, this study was able determine whether unique CAPs were detectable over multiple recording sessions. While most CAPs were only observed in single sessions, nearly one-third (29%) could be tracked in two or more consecutive recording days.

Average firing rates were tracked for each CAP at each level of anesthesia. In this study, baseline CAP activity was tracked with decreasing levels of isoflurane. Since nerve activity is suppressed at 2% isoflurane [35,45], levels used in this study were 1.5%, 1% and 0.5% as to not squash the neural response. CAPs with high firing rates at the highest level of anesthesia decreased with lowered anesthesia, as well as the inverse. Group 2 CAPs with FR_1.5_ ranging from 15-80 CAP/s did not vary in relation to isoflurane level, but the number of CAPs at the range disappeared as anesthesia was lowered. This group was nearly nonexistent in the awake animal recordings, while the high (>100) and low (<40) firing rate CAPs remained. There were CAPs with firing rates at the highest levels of anesthesia that match studies following baseline activity with increasing levels of anesthesia, with bursting related to breathing rates [14,18].

Of 575 extracted CAPs, 247 (43%) showed phase-locking preferences for cardiac, respiratory, or both signals. Strength of locking changed with level of anesthesia, but 90.7% of CAPs with some preference showed a FR_1.5_ between 0 and 15 CAP/s. The firing rates of CAPs with cardiac or respiratory phase-locking did not change with level of anesthesia, but those with phase-locking characteristics with both signals increased in firing rate at the lowest level of anesthesia. Similar relationships between vagal activity and other biological signals have been reported in other studies, in both preclinical and clinical models, including firing rate variations in relation to respiratory rate [18,28,46,47]. In a study inducing seizures in rats, the vagal neurogram was found to mirror heart rate [48], and intraneural electrodes in the human vagus nerve found multi-unit activity synced to the QRS interval with other tonically firing units locked with respiration or blood pressure fluctuations [29]. Because the VN has such mixed signaling, there is quite a bit of variation on reporting of firing rates and their preferential correlation with other biological signals, but this study shows that physiological-related signals can be recorded in a chronic murine model in a stable fashion. It will be crucial to classify currently uncategorized CAPs and signals, and their relationship with other organ functions, along with changes related to disease states.

Cytokine injections were performed in n = 2 animals, both three months after electrode implantation. Only one injection was done per week, and only one cytokine (IL-1β) was tested. In a total of six injection experiments, 7 CAPs responded to the cytokine injection with a change in firing rate and phase-locking characteristics. The response rate in this study was not as high as those published in previous work, as [15] reported up to 71.4% due to IL-1β injections, but those studies were done in an acute setting with an increased dosage of cytokine. The few animals tested in this work show that cytokine-specific sensory neural signals can be detected months after implantation, and future work can apply additional cytokines, dosages, and injection schedule while maintaining healthy and humane practices for managing these implanted mice.

There were a few other limitations in this work. Though the vagus nerve is a complicated network of neural efferents and afferents, this study only examined neural correlations with cardiac and respiratory signals, when there are so many other physiological biomarkers linked with the vagus nerve. Additionally, this work used extraneural electrodes that can suffer from low spatial resolution, compared to intraneural electrodes that may have higher SNRs and more spatially specificity. The vagus nerve may require more specific recording and stimulation methods to measure from and target organs, respectively. Lastly, there were difficulties with the surgery and implant, especially with securing the headcap and wires to prevent damage over time; five implants were terminated early due to mechanical failure by wire, electrode, or headcap damage.

This study establishes a novel, chronic recording model using an implanted cuff electrode to monitor CAPs from the murine VN for up to six months, showing long-term interface viability and stable SNRs. Unlike previous chronic studies in rats or previous acute, anesthetized studies in mice, this approach allows for longitudinal analysis in both awake and anesthetized mice, while tracking individual CAPs across multiple days, their firing rates and phase-locking characteristics with other physiological signals, as well as, in the awake case, movement using unsupervised machine learning models. These results reveal diverse CAP populations with varying degrees of physiological coupling. However, most recorded CAPs in this study remain unclassified, as well as their potential role in organ function. This chronic recording model provides a valuable platform in a flexible preclinical model to investigate how vagal activity may be modified based on disease severity, as well as provide a next step to develop closed-loop VNS by predicting flare-ups and tracking stimulation efficacy.

## Declaration of competing interests

The authors have no conflicts of interests to declare.

## Data Statement

Data and analysis code can be made available upon reasonable request to the corresponding authors.

## Funding

Funding for this research was provided by Northwell Health.

## Notes

### Competing Interest Statement

The authors have declared no competing interest.

