## Supplemental Figures for "Longitudinal characterization of compound action potentials in chronic vagus nerve recordings in mice"

Shubham Debnath et al.

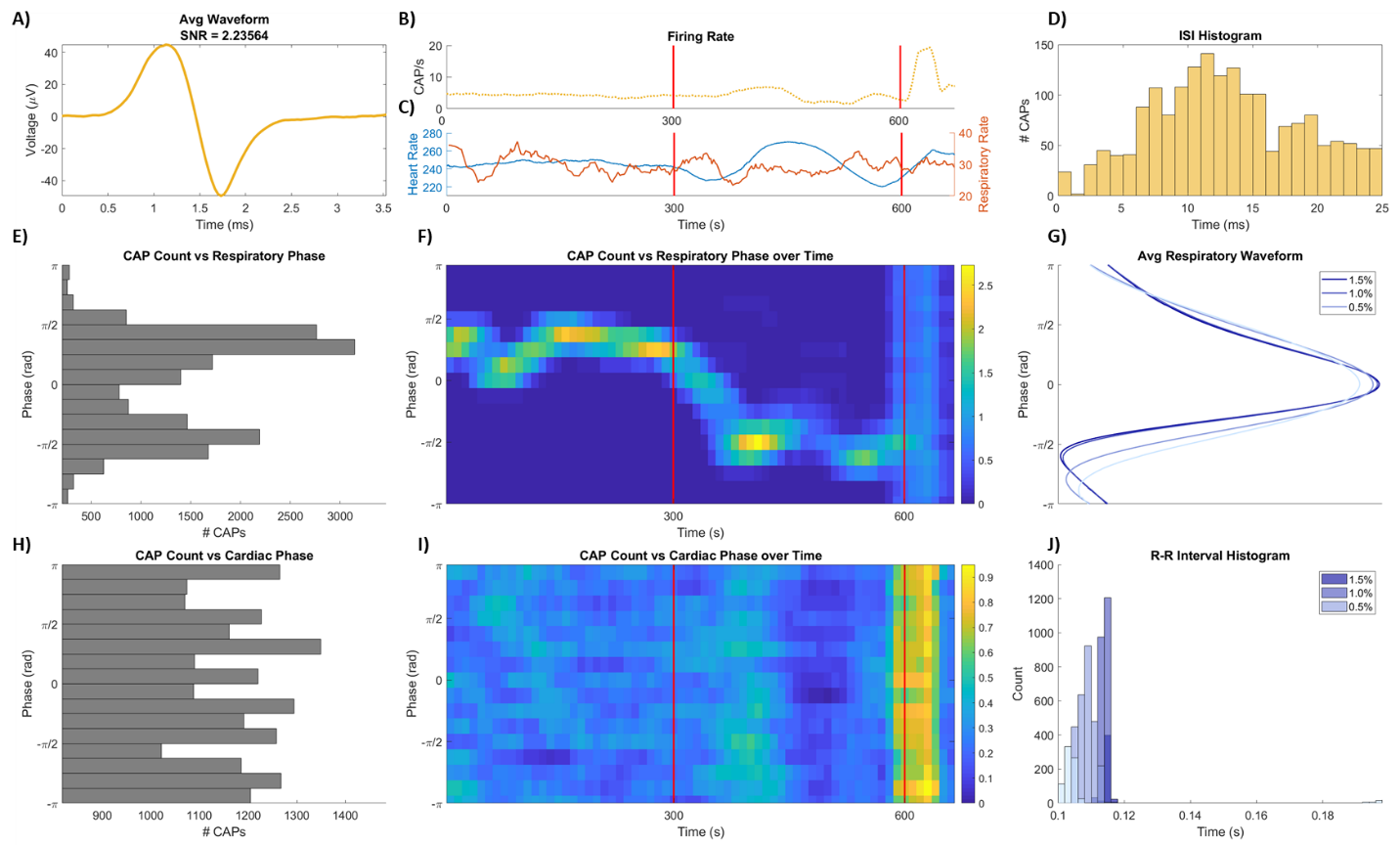
**Supplementary Figure 1. Example of CAP with phase-locking to cardiac signal under anesthesia.** Shown is an example of an extracted CAP and its features during baseline and after cytokine injection. Vertical red lines in the middle panels correspond to times of lowered anesthesia (from 1.5% to 1% then 1% to 0.5% isoflurane). (A) Average waveform and SNR. (B) Firing rate over the recording session, with (C) corresponding heart (blue) and respiratory rate (red) of the animal. (D) Inter-spike interval histogram. (E,F) CAP count vs respiratory phase, where 0 represents peak inhale, and –π and π represent halfway between the previous and next inhale, respectively. (G) Average shape of respiratory waveform at each level of anesthesia. (H,I) CAP count vs cardiac phase, where 0 represents peak of the R complex, and –π and π represent halfway between the previous and next R complex, respectively. (J) Histogram of R-R intervals at each level of anesthesia.

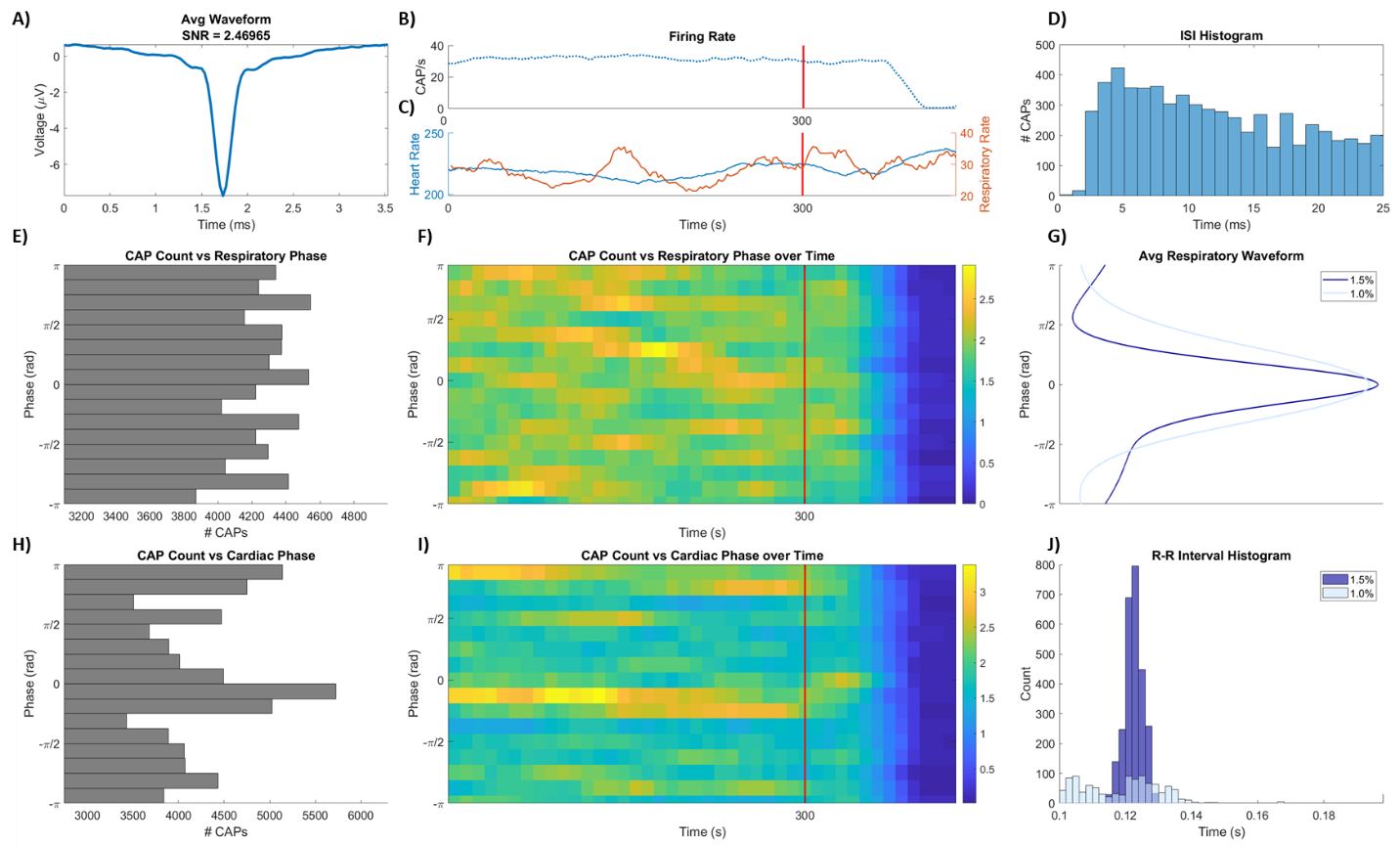
**Supplementary Figure 2. Example of CAP with phase-locking to respiratory signal under anesthesia.** Shown is an example of an extracted CAP and its features during baseline and after cytokine injection. Vertical red lines in the middle panels correspond to times of lowered anesthesia (from 1.5% to 1% isoflurane). (A) Average waveform and SNR. (B) Firing rate over the recording session, with (C) corresponding heart (blue) and respiratory rate (red) of the animal. (D) Inter-spike interval histogram. (E,F) CAP count vs respiratory phase, where 0 represents peak inhale, and –π and π represent halfway between the previous and next inhale, respectively. (G) Average shape of respiratory waveform at each level of anesthesia. (H,I) CAP count vs cardiac phase, where 0 represents peak of the R complex, and –π and π represent halfway between the previous and next R complex, respectively. (J) Histogram of R-R intervals at each level of anesthesia.

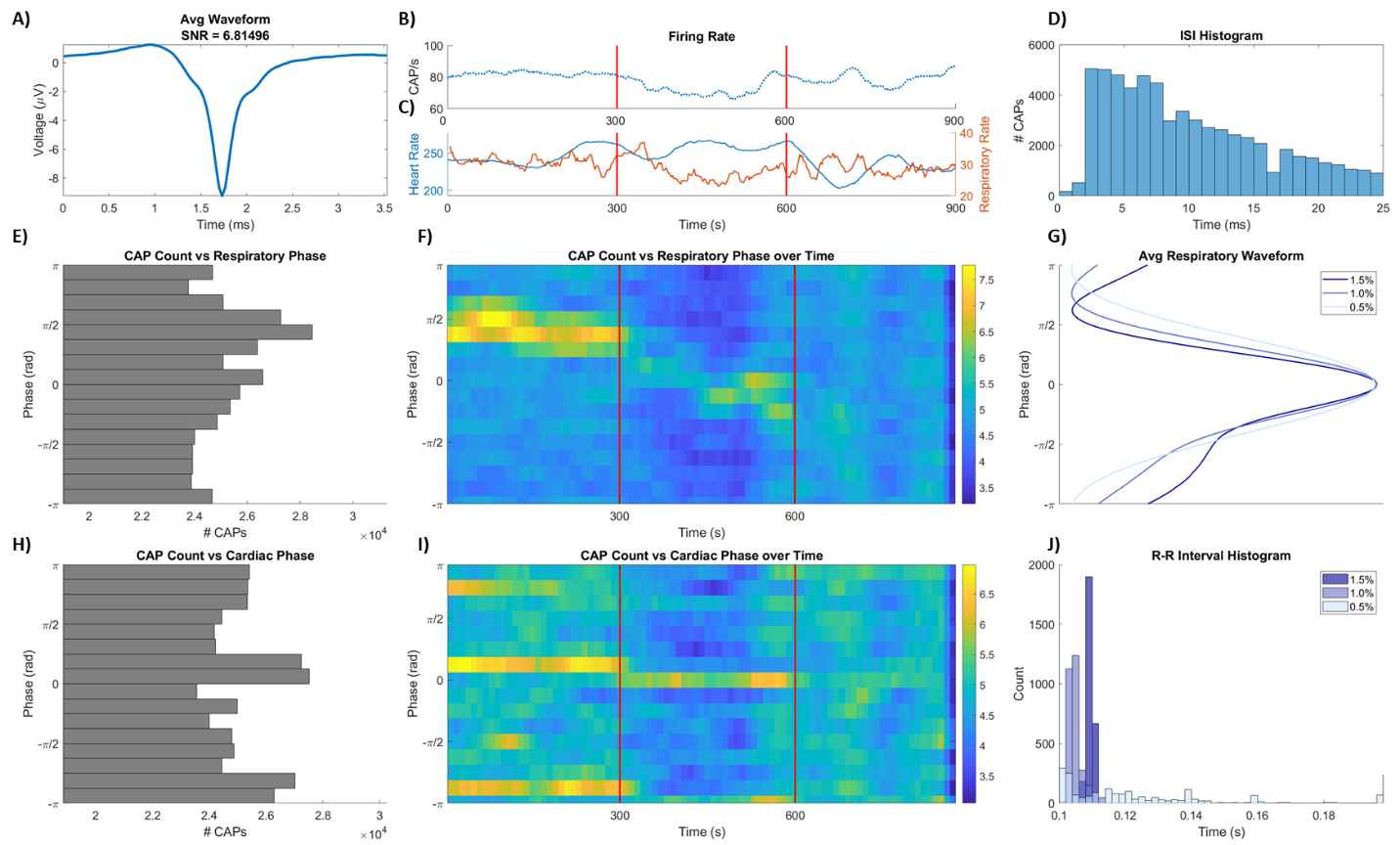
**Supplementary Figure 3. Example of CAP with phase-locking to both cardiac and respiratory signals under anesthesia.** Shown is an example of an extracted CAP and its features during baseline and after cytokine injection. Vertical red lines in the middle panels correspond to times of lowered anesthesia (from 1.5% to 1% then 1% to 0.5% isoflurane). (A) Average waveform and SNR. (B) Firing rate over the recording session, with (C) corresponding heart (blue) and respiratory rate (red) of the animal. (D) Inter-spike interval histogram. (E,F) CAP count vs respiratory phase, where 0 represents peak inhale, and –π and π represent halfway between the previous and next inhale, respectively. (G) Average shape of respiratory waveform at each level of anesthesia. (H,I) CAP count vs cardiac phase, where 0 represents peak of the R complex, and –π and π represent halfway between the previous and next R complex, respectively. (J) Histogram of R-R intervals at each level of anesthesia.
